# Quantitative investigation of factors relevant to the T cell spot test for tuberculosis infection in active tuberculosis

**DOI:** 10.1101/561886

**Authors:** Kui Li, Caiyong Yang, Zicheng Jiang, Shengxi Liu, Jun Liu, Chuanqi Fan, Tao Li, Xuemin Dong

## Abstract

**Background:** Previous qualitative studies suggested that the false negative rate of T cell spot test for tuberculosis infection (T-SPOT.TB) is associated with many risk factors in tuberculosis patients; However, more precise quantitative studies are not well known.

**Objective:** To investigate the factors affecting quantified T-SPOT.TB in patients with active tuberculosis.

**Methods:** We retrospectively analyzed the data of 360 patients who met the inclusion criteria. Using the levels of early secreted antigenic target 6 kDa (ESAT-6) and culture filtrate protein 10 kDa (CFP-10) as dependent variables, variables with statistical significance in the univariate analysis were subjected to optimal scaling regression analysis.

**Results:** The results showed that the ESAT-6 regression model had statistical significance (*P*-trend < 0.001) and that previously treated cases, CD4+ and platelet count were its independent risk factors (all *P*-trend < 0.05); their importance levels were 0.095, 0.596 and 0.100, respectively, with a total of 0.791. The CFP-10 regression model also had statistical significance (P-trend < 0.001); platelet distribution width and alpha-2 globulin were its independent risk factors (all *P*-trend < 0.05), their importance levels were 0.287 and 0.247, respectively, with a total of 0.534. The quantification graph showed that quantified T-SPOT.TB levels had a linear correlation with risk factors.

**Conclusion:** The test results of T-SPOT.TB should be given more precise explanations, especially in patients with low levels of CD4+, platelet, alpha-2 globulin and high platelet distribution width.

## Introduction

The interferon-gamma release assay (IGRA) represents one of the most important advances in the immunodiagnosis of *Mycobacterium tuberculosis* (MTB) infection in the past two decades. As a new adjuvant method for the diagnosis of MTB infection, IGRA has been widely applied and accepted clinically. In principle, IGRA determines whether the subject is infected with MTB by examination of the levels of released γ-interferon (IFN-γ) after stimulation of whole blood or peripheral blood mononuclear cells (PBMCs) with MTB-specific antigen. This test is not affected by Bacillus Calmette-Guerin (BCG) vaccination [1], a feature that is very beneficial in countries such as China in which general BCG vaccination is practiced. Currently, the T cell spot test for tuberculosis infection (T-SPOT.*TB*) is the main IGRA test method; it provides intuitive and reproducible results and quantitatively reflects the number of IFN-γ secreting cells in preparations of PBMCs [2].

IGRA still has a certain false negative rate among patients with tuberculosis (TB). Previous studies reported that negative bacteria in sputum [3-5], hypoproteinemia [6-8], combined HIV infection [4, 7, 9], anti-TB treatment [10, 11], medical history [8, 12], anemia [6, 13], diabetes [14], parasitic infections [13], noncavitary lesions in the lung [5], fall and winter seasons [15], increased human leukocyte antigen DRB1-0701 allele [16] and non-Hispanic white or Asian ethnicity [6, 9] are risk factors for false negative IGRA. The association between IGRA and age [3, 7-9, 12, 13, 15-18], body mass index [12, 16] and reduced lymphocyte levels [3, 5, 7, 8, 17, 19, 20] is inconsistent among previous studies. Some of these studies were qualitative studies with small samples and did not consider the possibility that different antigen risk factors might have confounded or biased the results. In addition, there is a lack of quantitative examination of the association between spot-forming cells (SFCs) within PBMCs and clinical and laboratory characteristics. As a result, further research is urgently needed. Our main objective based on population analysis is to identify factors associated with different antigen quantification results of T-SPOT.*TB* in active TB (ATB), with the goal of providing more reference evidence for the accurate application of T-SPOT.*TB*.

## Methods and materials

### Study populations

We retrospectively analyzed 360 pathogenically positive TB patients who were hospitalized in Ankang Central Hospital, China between October 1, 2016 and June 30, 2018. We collected data on 29 variables associated with five aspects including the patients’ general status, bacteriology, imaging, routine examination, protein electrophoresis and immunology. The subject screening process is illustrated in (Fig 1).

**Fig 1.**
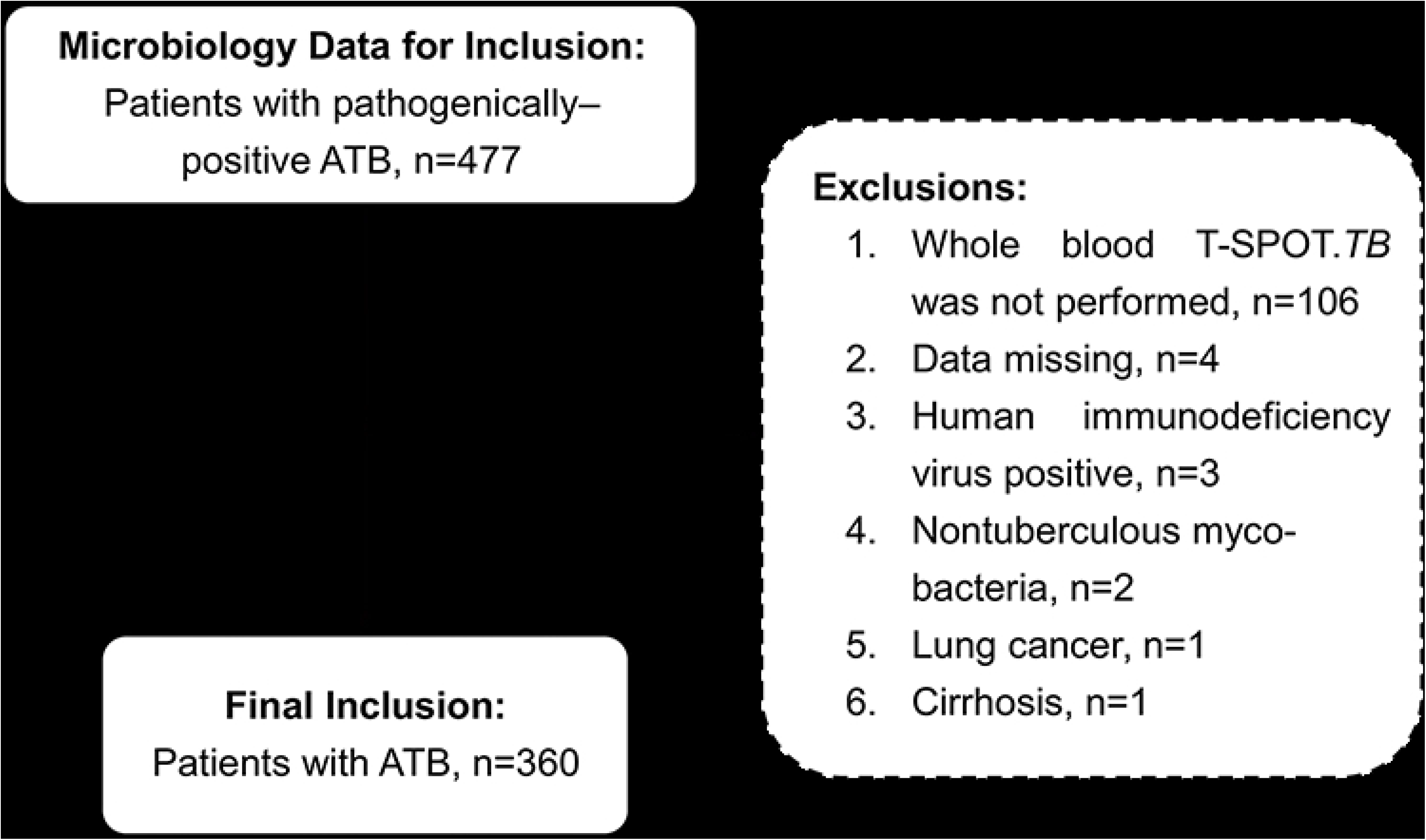
Description of the sample. ATB, active tuberculosis.

This study was approved by the ethics committee of Ankang Central Hospital. Informed consent was not required because the study did not put the patients at risk. Prior to analysis, the patients’ identities were protected by anonymity and by the use of codes.

### Inclusion and exclusion criteria

The inclusion criteria were as follows [21]: (1) The MTB nucleic acid test was positive, and patients had suspected symptoms of TB. (2) Patients had one sputum smear that was positive for acid-fast staining or positive sputum culture, and thoracic imaging showed lesions conforming to ATB and/or suspected symptoms of TB. (3) Patients were subjected to the T-SPOT.*TB* test, and the results were accessible.

The exclusion criteria were as follows: (1) Patients who were taking immunosuppressants. (2) Patients who did not undergo IGRA examination. (3) Patients who had positive HIV antibody. (4) Patients with nontuberculous mycobacteria infection. (5) Patients with combined cirrhosis. (6) Patients with combined tumor.

### T-SPOT.TB assay

The T-SPOT.*TB* diagnosis kit was provided by Shanghai Fosun Long March Medical Science Co., LTD (Oxford Immunotec Ltd., United Kingdom). Specimens were examined within 2 h of collection at room temperature (18-25 °C) in a sterile environment. (1) RPMI 1640 medium was mixed with an equal volume of a whole blood sample. (2) Three milliliters of Ficoll human lymphocyte isolation solution was added to the first centrifuge tube; 6-9 ml of diluted whole blood was then slowly added to the solution, and the tube was centrifuged at 1000 × g for 22 min. (3) Approximately 5 ml of the layer containing PBMCs was transferred to a second centrifuge tube, and the mixture was brought to a volume of 10 ml with RPMI 1640; the tube was then inverted and centrifuged at 600 × g for 7 min. (4) The supernatant was discarded, the pellet was resuspended in 1 ml of medium, and RPMI 1640 was added to bring the final volume to 10 ml. After mixing by inverting the tube, the sample was centrifuged at 350 × g for 7 min. (5) The supernatant was discarded, and the pellet was resuspended in AIM-V medium at a density of 2.5 × 10^6^ cells/ml. (6) Fifty microliters of negative control (ALM-V), antigen A (early secreted antigenic target 6 kDa, ESAT-6), antigen B (culture filtrate protein 10 kDa, CFP-10) and positive control (PHA) were added sequentially to an IFN-γ antibody-coated microplate; 100 µl of diluted cells were then added to each well, and the plate was placed in a CO2 incubator (37 °C, 5% CO2) for 20 hr. (7) After removal of the medium, each well was washed four times with 200 µl of PBS and incubated with 50 µl of secondary antibody working solution (1:200) for 1 h at 2-8 °C. (8) After removal of the secondary antibody, each well was washed four times with PBS, for four times and added 50 µl of substrate chromogen solution was added to each well, and the plate was allowed to stand for 10 min in the dark. The reaction was terminated by washing the plate with distilled water. (9) SFCs were automatically counted using a plate reader (ES-15, Shanghai Fosun Long March Medical Science Co., Ltd., Shanghai, China). When the counts could not be determined using the plate reader, the SFCs were manually counted using a microscope. The results were recorded and interpreted as recommended by the kit manufacturer.

### Measurement of TB

Sputum or body fluid specimens were first examined using Auramine O fluorescence staining (KRJ/TTR500 automatic smear staining machine, Xiangyang Courager Medical Apparatus, Xiangyang, China) and were then subjected to MTB nucleic acid amplification (RNA constant temperature amplification, Roche LightCycler 480 II Real-time PCR Cycler; Rendu Biotechnology [Shanghai, China]; PCR-fluorescence method, Applied Biosystems 7500 Real-Time PCR System; DAAN Gene of Sun Yat-sen University [Guangzhou, China]) and rapid drug tolerance gene analysis (DNA microarray chip method; CapitalBio Corporation, Chengdu, China). Finally, the specimens were subjected to MTB culture (Roche fixed culture test), bacterial species identification and drug sensitivity testing (Ratio methods, BaSo Diagnostics INC., ZHUHAI). Using the Roche solid culture as the standard and no repeated counting, bacterial load was reported in accordance with the “*Diagnostic Criteria and Principles of Management of Infectious Pulmonary Tuberculosis*” [21] (S1 Table). The laboratory meets the national P3 requirements and accepts the quality control and management of the national reference laboratory. All operators received special training in the testing methods and in operation of the apparatus.

### Measurement of other variables

Body mass index was calculated by dividing the individual’s body weight (kg) by the square of his or her height (m). Body weight and height were measured at the time of admission. Behavioral risk factors (such as smoking and exposure to dust) and medical history were collected by physicians. The diagnostic criteria for diabetes were random blood glucose ≥11.1 mmol/L or fasting blood glucose ≥7.0 mmol/L [22]. The severity of lung imaging was graded according to the criteria set by the National Tuberculosis Association of America [23] (S2 Table). Whole blood cell counts were determined using a Sysmex XN-9000 automatic blood fluid analyzer and supporting reagents. Serum protein electrophoresis was performed with the Sebia Capillarys 2 Flex Piercing detection platform and supporting reagents (Pare Technologique Leonard de Vinci, CP8010-Lisses 91008 EVRY CEDEX, France). Lymphocyte subsets were detected using the Mindray flow cytometer BriCyte E6 and a kit from BD.

### Statistical analysis

The data were analyzed using SPSS 22.0 software (IBM SPSS Statistics). Non-normally distributed data were expressed as median and interquartile range. Wilcoxon’s rank sum test was used to compare two-sample dichotomous variables. The rank data and measurement data were analyzed using the Spearman relationship test, and the statistically significant variables were subjected to optimal scaling regression analysis. The missing values of the independent variables were replaced by the series mean. Adobe Photoshop CS6 software (version 13.0×32) was used for graphing. *P*-trend < 0.05 was considered statistically significant.

## Results

Univariate analysis showed: There was statistical significance in the yes or on of previously treated cases with ESAT-6 (*P*-trend < 0.05). CD4+, CD8+, platelet (PLT), prealbumin, albumin, albumin/globulin and beta-1 globulin were positively associated with ESAT-6 (all *P*-trend < 0.05). Alpha-1 globulin was negatively associated with ESAT-6 (*P*-trend < 0.05). CD4+, CD8+, PLT, prealbumin, albumin and alpha-2 globulin were positively associated with CFP-10 (all *P*-trend < 0.05). Age and platelet distribution width (PDW) were negatively associated with CFP-10 (all *P*-trend < 0.05). The associations with other independent variables, including gender, sputum bacterial load and body mass index etc, were not statistically significant (all *P*-trend > 0.05) (Tables 1-3).

**Table 1.**
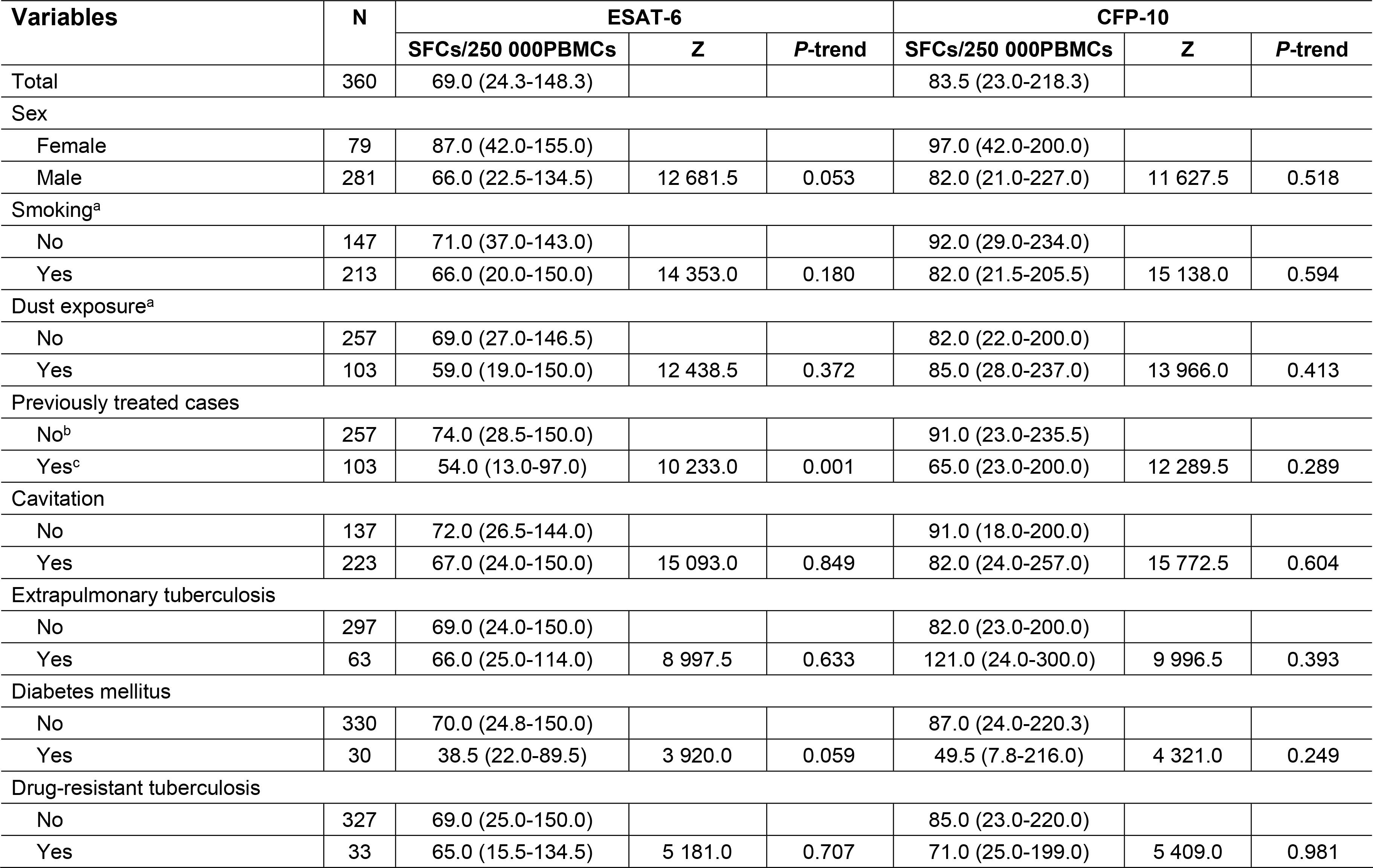
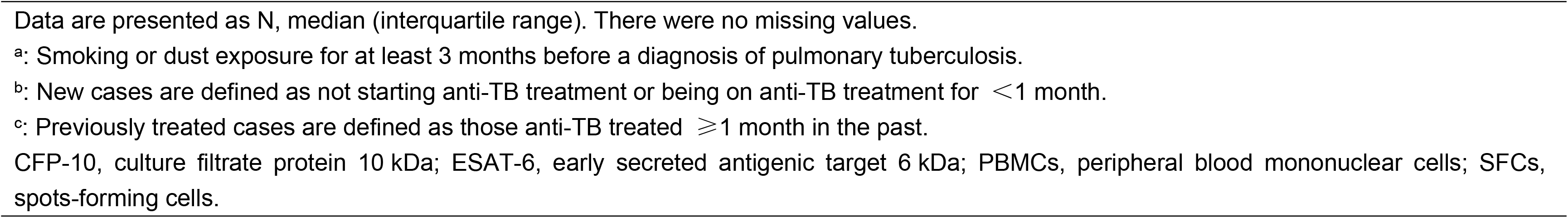
Univariate analysis of T-SPOT.TB (nominal variables).

**Table 2.** Univariate analysis of T-SPOT.TB (ordinal variables).

| Variables | N | ESAT-6 |  |  | CFP-10 |  |  |
| --- | --- | --- | --- | --- | --- | --- | --- |
|  |  | SFCs/250 000PBMCs | SRCC | P-trend | SFCs/250 000PBMCs | SRCC | P-trend |
| Stage<sup>a</sup> |  |  |  |  |  |  |  |
| Minimal/Mild | 2 | 77.5 (0.0-77.50) |  |  | 85.5 (22.0-85.5) |  |  |
| Moderate | 143 | 78.0 (38.0-150.0) |  |  | 97.0 (41.0-264.0) |  |  |
| Advanced | 215 | 64.0 (22.0-126.0) | -0.095 | 0.071 | 82.0 (15.0-211.0) | 0.000 | 0.995 |
| Grade<sup>b</sup> |  |  |  |  |  |  |  |
| DNA/RNA positive | 80 | 68.0 (27.3-99.5) |  |  | 84.0 (22.5-251.8) |  |  |
| The number of colony | 24 | 104.0 (51.8-203.5) |  |  | 132.5 (24.8-262.5) |  |  |
| 1+ | 102 | 67.0 (21.8-150.0) |  |  | 77.0 (22.8-205.0) |  |  |
| 2+ | 50 | 59.5 (25.5-142.3) |  |  | 81.0 (21.3-291.8) |  |  |
| 3+ | 70 | 69.0 (24.5-150.0) |  |  | 86.0 (26.8-200.0) |  |  |
| 4+ | 34 | 65.5 (30.3-126.8) | -0.075 | 0.158 | 59.5 (19.3-154.8) | -0.029 | 0.587 |

Data are presented as N, median (interquartile range). There were no missing values.
<sup>a</sup>: The staging standard of National Tuberculosis Association of the United States of America.
<sup>b</sup>: Smear grading before treatment.
CFP-10, culture filtrate protein 10 kDa; DNA, deoxyribonucleic acid; ESAT-6, early secreted antigenic target 6 kDa; RNA, ribonucleic acid; SRCC, spearman`s rho correlation coefficient.

**Table 3.** . Univariate analysis of T-SPOT.*TB* (numeric variables).

| Variables | Missing Data |  | Median (interquartile range)<sup>a</sup> | ESAT-6 |  | CFP-10 |  |
| --- | --- | --- | --- | --- | --- | --- | --- |
|  | N | (%) |  | SRCC | P-trend | SRCC | P-trend |
| Age, years | 0 | (00.0) | 53.0 (36.0-66.0) | -0.102 | 0.054 | -0.107<sup>*</sup> | 0.042 |
| Body-mass index, kg/m<sup>2</sup> | 6 | (1.7) | 19.1 (17.6-21.2) | 0.030 | 0.565 | 0.013 | 0.801 |
| Duration of symptoms, days | 0 | (00.0) | 90.0 (30.0-496.3) | -0.073 | 0.165 | 0.025 | 0.630 |
| Hemoglobin, g/L | 0 | (00.0) | 118.0 (105.0-134.8) | 0.034 | 0.518 | 0.016 | 0.765 |
| Platelet, ×10<sup>9</sup>/L, | 0 | (00.0) | 258.0 (191.0-320.8) | 0.173<sup>**</sup> | 0.001 | 0.186<sup>**</sup> | < 0.001 |
| Platelet distribution width, fL | 0 | (00.0) | 14.0 (11.7-16.2) | -0.090 | 0.089 | -0.116<sup>*</sup> | 0.027 |
| Erythrocyte sedimentation rate, mm/H | 32 | (8.9) | 49.9 (28.3-69.0) | -0.070 | 0.184 | -0.028 | 0.594 |
| Prealbumin, mg/L | 39 | (10.8) | 143.6 (84.3-184.8) | 0.187<sup>**</sup> | < 0.001 | 0.104<sup>*</sup> | 0.050 |
| Albumin, g/L | 1 | (0.3) | 31.9 (27.6-36.5) | 0.159<sup>**</sup> | 0.002 | 0.108<sup>*</sup> | 0.040 |
| Globulin, g/L | 5 | (1.4) | 32.6 (28.9-36.1) | 0.014 | 0.795 | 0.049 | 0.356 |
| Albumin to globulin ratio | 5 | (1.4) | 1.0 (0.8-1.2) | 0.105<sup>*</sup> | 0.047 | 0.054 | 0.303 |
| Alpha-1 globulin, g/L | 40 | (11.1) | 4.9 (3.7-5.6) | -0.122<sup>*</sup> | 0.020 | -0.077 | 0.146 |
| Alpha-2 globulin, g/L | 40 | (11.1) | 7.6 (6.5-8.6) | 0.091 | 0.085 | 0.141<sup>**</sup> | 0.007 |
| Beta-1 globulin, g/L | 40 | (11.1) | 3.7 (3.3-4.1) | 0.111<sup>*</sup> | 0.036 | 0.058 | 0.270 |
| Beta-2 globulin, g/L | 40 | (11.1) | 3.7 (3.2-4.1) | 0.034 | 0.514 | 0.063 | 0.233 |
| Gamma globulin, g/L | 40 | (11.1) | 13.0 (10.4-14.7) | 0.012 | 0.823 | 0.014 | 0.797 |
| CD4<sup>+</sup> T lymphocytes, cells/μL | 89 | (24.7) | 302.0 (203.0-445.0) | 0.391<sup>**</sup> | < 0.001 | 0.230<sup>**</sup> | < 0.001 |
| CD8<sup>+</sup> T lymphocytes, cells/μL | 89 | (24.7) | 253.0 (164.0-379.0) | 0.291** | < 0.001 | 0.190** | < 0.001 |
| CD4<sup>+</sup> to CD8<sup>+</sup> ratio | 89 | (24.7) | 1.2 (0.9-1.7) | 0.049 | 0.354 | -0.042 | 0.425 |

<sup>a</sup>: Data are presented as the value of variable.
\*: Correlation is significant at the 0.05 level (2-tailed).
\*\*: Correlation is significant at the 0.01 level (2-tailed).
CFP-10, culture filtrate protein 10 kDa; ESAT-6, early secreted antigenic target 6 kDa; SRCC, spearman`s rho correlation coefficient.

Optimal scaling regression analysis showed that the regression models were statistically significant (ESAT-6, adjusted R-squared = 0.147, F = 7.884, *P*-trend < 0.001; CFP-10, adjusted R-squared = 0.061, F = 3.890, *P*-trend < 0.001). Analysis of all the included factors showed that previously treated cases, CD4+ and PLT were significantly associated with ESAT-6; their importance levels were 0.095, 0.596 and 0.100, respectively, with a total of 0.791. PDW and alpha-2 globulin were also significantly associated with CFP-10; their importance levels were 0.287 and 0.247, respectively, with a total of 0.534. The tolerances of the included factors in the two models were > 0.1, indicating that there was no serious collinearity among the factors and that the regression models were reliable (Table 4).

**Table 4.** Optimal scale regression analysis of T-SPOT.TB.

| Variables | Beta | Standardized coefficients<br>Std.Error | df | F | Sig. | Tolerance |  | Importance |
| --- | --- | --- | --- | --- | --- | --- | --- | --- |
|  |  |  |  |  |  | After transformation | Before transformation |  |
| Early secreted antigenic target 6<sup>a</sup> |  |  |  |  |  |  |  |  |
| Previously treated cases | 0.103 | 0.046 | 1 | 5.110 | 0.024 | 0.948 | 0.948 | 0.095 |
| Platelet, ×10<sup>9</sup>/L, | 0.115 | 0.054 | 1 | 4.471 | 0.035 | 0.818 | 0.818 | 0.100 |
| Prealbumin, mg/L | 0.043 | 0.072 | 1 | 0.366 | 0.545 | 0.492 | 0.492 | 0.040 |
| Albumin, g/L | 0.080 | 0.094 | 1 | 0.738 | 0.391 | 0.253 | 0.253 | 0.076 |
| Albumin to globulin ratio | -0.075 | 0.082 | 1 | 0.819 | 0.366 | 0.274 | 0.274 | -0.039 |
| Alpha-1 globulin, g/L | -0.092 | 0.056 | 1 | 2.630 | 0.106 | 0.725 | 0.725 | 0.072 |
| Beta-1 globulin, g/L | -0.016 | 0.062 | 1 | 0.067 | 0.796 | 0.617 | 0.617 | -0.008 |
| CD4<sup>+</sup> T lymphocytes, cells/μL | 0.277 | 0.075 | 1 | 13.539 | 0.000 | 0.598 | 0.598 | 0.596 |
| CD8<sup>+</sup> T lymphocytes, cells/μL | 0.048 | 0.063 | 1 | 0.588 | 0.444 | 0.658 | 0.658 | 0.069 |
| Culture filtrate protein 10<sup>b</sup> |  |  |  |  |  |  |  |  |
| Age, years | -0.022 | 0.057 | 1 | 0.153 | 0.696 | 0.867 | 0.867 | 0.018 |
| Platelet, ×10<sup>9</sup>/L, | 0.027 | 0.068 | 1 | 0.161 | 0.688 | 0.634 | 0.634 | 0.047 |
| Platelet distribution width, fL | -0.149 | 0.058 | 1 | 6.549 | 0.011 | 0.701 | 0.701 | 0.287 |
| Prealbumin, mg/L | -0.032 | 0.076 | 1 | 0.179 | 0.672 | 0.539 | 0.539 | -0.021 |
| Albumin, g/L | 0.118 | 0.070 | 1 | 2.830 | 0.093 | 0.488 | 0.488 | 0.131 |
| Alpha-2 globulin, g/L | 0.133 | 0.049 | 1 | 7.464 | 0.007 | 0.917 | 0.917 | 0.247 |
| CD4<sup>+</sup> T lymphocytes, cells/μL | 0.115 | 0.071 | 1 | 2.648 | 0.105 | 0.608 | 0.608 | 0.236 |
| CD8<sup>+</sup> T lymphocytes, cells/μL | 0.031 | 0.063 | 1 | 0.245 | 0.621 | 0.652 | 0.652 | 0.055 |

<sup>a</sup>: R Square = 0.169; Adjusted R Square = 0.147; F = 7.884; *P*-trend < 0.001.
<sup>b</sup>: R Square = 0.081; Adjusted R Square = 0.061; F = 3.890; *P*-trend < 0.001.

The quantification graph after variable conversion showed that (1) the quantified ESAT-6 level of previously treated cases was lower than new cases (Fig 2A); (2) the quantified ESAT-6 level increased as CD4+ and PLT increased (Figs 2B and 2C); (3) the quantified CFP-10 level increased as PDW and alpha-2 globulin increased (Figs 2D and 2E).

**Fig 2.**
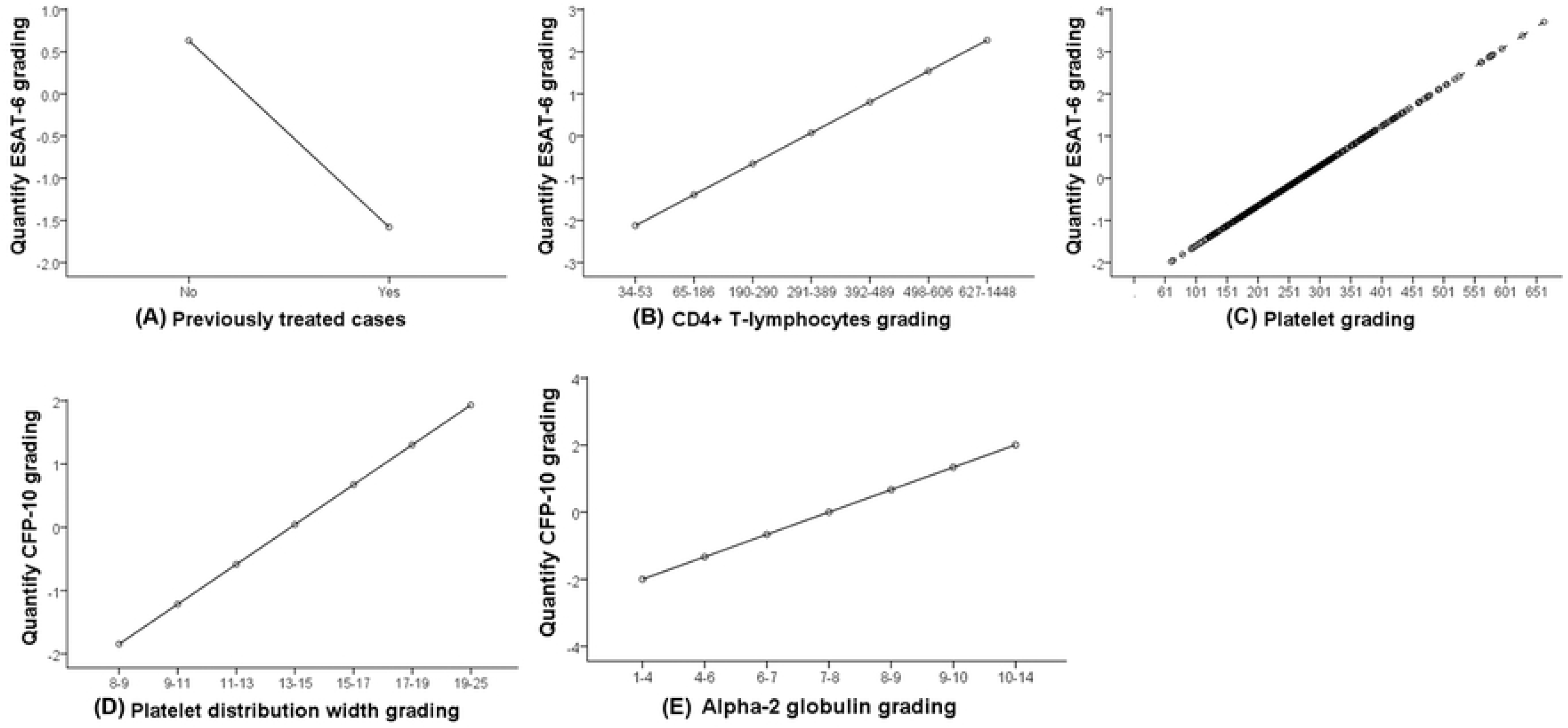
Conversion graph between the quantified values of the independent and dependent variables. (A) Patients with a history of previous treatment showed decreased ESAT-6 grading. (B) Quantified ESAT-6 grading increased with an increase in the number of CD4+ T lymphocytes. (C) Quantified ESAT-6 grading increased with an increase in the number of PLT. (D) Quantified CFP-10 grading increased with an increase in PDW grading. (E) Quantified CFP-10 grading increased with an increase in alpha-2 globulin grading.

## Discussion

In this study, the false-negative rates of ESAT-6, CFP-10 and T-SPOT.*TB* were 8.61% (31/360), 10.00 (36/360) and 3.33% (12/360), respectively, based on the recommended cut-off value (<6 SFCs/250,000 PBMCs). The observed false-negative rate of T-SPOT.*TB* is lower than the T-SPOT.*TB* false-negative rate of 6.74% (61/905) reported in a previous study [24]. This discrepancy may be due to inconsistencies in case inclusion criteria and to the existence of different comorbidities.

This study showed that the ESAT-6 of previously treated cases was significantly lower than new cases. The causes may include the following: (1) As the disease progresses, the cellular immune function of the body further declines, and the corresponding immune reactions are gradually weakened. Cell activation or immunosuppression is further aggravated, and this aggravation weakens the immune response of the T lymphocytes to MTB and leads to a reduced number of cells releasing IFN-γ. (2) MTB may directly and continuously downregulate helper T cells (Th1), leading to reduced function of Th1 in secreting IFN-γ [25]. (3) Following anti-TB treatment, the MTB is gradually cleared from the body. However, the TB antigen-specific effector T cells are also reduced in number or even disappear, which may lead to a decline in ESAT-6. This outcome is consistent with a previous study [26].

The role of PLT is not limited to acute hemostasis and vascular wall repair. Increasing evidence indicates that PLT are the key effector cells through which the host regulates inflammatory responses and that PLT contribute to the initiation and spreading of local and systemic inflammation [27, 28]. PLT may induce T lymphocyte adhesion by secreting the key chemical inducer RANTES (regulated on activation, normal T-cell expressed and secreted, also known as CCL5), thereby regulating the functions of T lymphocytes [27, 29, 30]. This study found that changes in the number of PLT were positively associated with ESAT-6 secretion, an effect that may be due to an overall increase in IFN-γ through indirect regulation of T lymphocytes. Although the detailed pathway is still unclear, our finding is consistent with the study of Kim et al. [31] in which it was shown that the false-positive rate was associated with low levels of PLT. In multivariate analysis, we found no association between PLT and CFP-10.

When MTB infects the body, MTB sensitizes T lymphocytes after presentation by antigen-presenting cells. T lymphocytes rapidly proliferate and differentiate into functional effector T cells. Therefore, the number of lymphocytes, especially the number and function of CD4+ T lymphocytes, may affect the results of the T-SPOT.*TB* test. During the T-SPOT.*TB* test, the number of cells in the sample was standardized by adjusting the cell concentration to a standard value; to some extent, this standardization decreased the effect of cell number on the test results. However, in the current study, we found that CD4+ and CD8+ counts were associated with T-SPOT.*TB* in the univariate analysis and that in the multivariate analysis only CD4+ count was associated with ESAT-6. The quantification graph following variable conversion showed that increased CD4+ counts were associated with higher ESAT-6 values, and the importance level was relatively high, suggesting that CD4+ count is a main factor affecting ESAT-6. CD8+ count is not associated with T-SPOT.*TB*, CD4+ or CFP-10, suggesting that CD4+ cells may play a more important role in MTB infection than CD8+ cells [32]. This finding is inconsistent with a previous study in which it was shown that the immune status of the subject had minimal effect on IGRA-ELISPOT test results [33] but is consistent with the study of Zhang et al. [5]. This may be related to one or more of the following factors. (1) When a patient infected with TB is immunized, large numbers of lymphocytes accumulate at the lesion, leading to a decrease in the number of lymphocytes in the peripheral circulation and a reduction in the number of active T lymphocytes; this eventually leads to reduced numbers of SFCs in the T-SPOT.*TB* test. (2) ESAT-6 is more closely associated with the transcription and expression of IFN-γ by T lymphocytes [34]. (3) During immunization, CD4+ T cells preferentially release interferon, whereas CD8+ T cells preferentially lyse antigen-presenting cells [35]. Further study and analysis are required to determine the importance of changes in the number of CD4+ T lymphocytes in this process.

PDW is a parameter that reflects the variation in PLT volume. Approximately 50-80% of PLT volume is contributed by α-granules, and the various cytokines secreted in α-granules participate in important pathways associated with PLT immune regulation [36, 37]. P-selectin is a key component of α-granules, P-selectin deficiency leads to higher bacterial load in the lungs [38]. It is possible that when the body is stimulated by the MTB antigen, the number of PLT with small volumes and lacking P-selectin, leading to an overall increase in CFP-10 through cytokine regulation. CFP-10 may indirectly reflect potential markers of PLT function through α-granules. To date, there have been no relevant studies on the detailed regulation of this pathway.

In an early study of patients with TB, alpha-2 globulin and γ-globulin were elevated, alpha-2 globulin elevation was associated with exudative lesions, and a reduced albumin/alpha-2 globulin ratio indicated a poor prognosis [39]. A recent study showed that failure of serum globulin to recover by two months after treatment was associated with a need for extended treatment [40]. These studies showed that globulin was altered in patients with TB. In this study, we found that 29.38% (94/320) of patients had alpha-2 globulin levels that were greater than the upper limit of the normal range (8.5 g/L). Univariate screening showed that albumin was associated with T-SPOT.*TB*, consistent with previous studies [6-8, 31]. However, multivariate analysis showed that alpha-2 globulin is another factor that is associated with changes in CFP-10 levels, which in turn may be associated with globulin elevation and peripheral T cell reactions (unpublished data), including the production of MTB-specific IFN-γ. However, the specific pathway of this effect is unclear, and there are no similar previous studies.

The limitations of this study are as follows. First, the amount of information available for interpreting T-SPOT.*TB* variation was low, and other unknown factors remain to be investigated. Second, unlike demographic, clinical and laboratory characteristics, the genetic causes of false-negative T-SPOT.*TB* results cannot be studied. Finally, the current data do not explain the different impact of the same variable on ESAT-6 and CFP-10. However, this is the first study to demonstrate the effects of PDW and alpha-2 globulin on CFP-10 and is the largest study to date to quantify the association of different antigens with T-SPOT.*TB* in ATB.

In T-SPOT.TB-assisted diagnosis of patients with ATB, special attention should be given to the influence of previously treated cases and number of CD4+ cells and PLT on ESAT-6 and to the influence of PDW and alpha-2 globulin on CFP-10. Detailed assessment of these factors may help us accurately understand the diagnosis of TB using T-SPOT.*TB*.

## Supporting information

**<u>S1 Table</u>.** Smear grading report standard (DOC).

**<u>S2 Table</u>.** Thoracic CT scan image classification criteria(DOC).

## Acknowledgments

American Journal Experts (Durham, North Carolina) provided assistance in translating the manuscript.

## Author Contributions

**Conceptualization:** Kui Li, Caiyong Yang, Zicheng Jiang.

**Data curation:** Kui Li, Shengxi Liu, Jun Liu, Chuanqi Fan, Xuemin Dong.

**Formal analysis:** Kui Li.

**Investigation:** Kui Li, Shengxi Liu, Jun Liu, Chuanqi Fan, Tao Li, Xuemin Dong

**Methodology:** Kui Li, Caiyong Yang.

**Project administraton:** Kui Li, Zicheng Jiang.

**Resources:** Jun Liu, Xuemin Dong.

**Software:** Kui Li.

**Supervision:** Caiyong Yang.

**Validation:** Kui Li, Shengxi Liu.

**Visualization:** Kui Li, Chuanqi Fan.

**Writing – original draft:** Kui Li, Caiyong Yang.

**Writing – review & editing:** Kui Li, Caiyong Yang, Zicheng Jiang, Shengxi Liu, Jun Liu, Chuanqi Fan, Tao Li, Xuemin Dong.

