## Supplemental Table 1 for "Quantitative investigation of factors relevant to the T cell spot test for tuberculosis infection in active tuberculosis"

**S1 Table. Smear grading report standard**^a^.

**Sputum smear**

+: 1–9 bacteria/50 fields;

1+: 10–49 bacteria/50 fields;

2+: 1–9 bacteria/field;

3+: 10–90 bacteria/field;

4+: ≥ 100 bacteria/field.

At least 50 fields were observed for the 2+ reports and at least 20 fields were observed for 3+ and above results.

**Sputum culture**

+: the actual colony count was reported, as the bacterial colony growth was less than 1/4 of the slope surface area;

1+: bacterial colony growth accounted for 1/4 of the slope surface area;

2+: bacterial colony growth accounted for 1/2 of the slope surface area;

3+ bacterial colony growth accounted for 3/4 of the slope surface area;

4+: bacterial colony growth accounted for entire the slope surface area.

^a^: Refers to *Diagnostic Criteria and Principles of Management of Infectious Pulmonary Tuberculosis*（GB15987-1995）*.*
