## Supplemental Table 2 for "Quantitative investigation of factors relevant to the T cell spot test for tuberculosis infection in active tuberculosis"

**S2 Table.** Thoracic CT scan image classification criteria^a^.

Stage 1 (minimal/mild)

Mild to moderately dense lesions with no cavities and involving only part of one lung or both lungs. The entire range is smaller than the volume of the lung on one side above the junction between the second rib and the sternum.

Stage 2 (moderate)

Lesions were in one lung or both lungs, but the entire range did not exceed any of the following: (1) Small or moderate diffuse lesions with a distribution that did not exceed the entire area of one lung. If lesions were in both lungs, the total area of the lesions did not exceed the area of one lung. (2) Highly dense fusion lesions that did not exceed one-third of the volume of a single lung. (3). When there were cavities, the largest diameter of the cavity was less than 4 cm.

Stage 3 (advanced)

Cases in which the lesion range exceeded the range described above for moderate lesions.

^a^: National Tuberculosis Association of the USA: Diagnostic standards and classification of tuberculosis. 1961, National Tuberculosis Association, New York.
